# Osteoprotegerin expression in liver is induced by IL-13 through TGF-β

**DOI:** 10.1101/2020.12.11.421479

**Authors:** Adhyatmika Adhyatmika, Kurnia S. S. Putri, Emilia Gore, Keri A. Mangnus, Catharina Reker-Smit, Detlef Schuppan, Leonie Beljaars, Peter Olinga, Barbro N. Melgert

## Abstract

1.

**Backgrounds:** Osteoprotegerin (OPG) is a profibrotic mediator produced by myofibroblasts under influence of transforming growth factor β (TGFβ). Its expression in experimental models of liver fibrosis correlates well with disease severity and treatment responses. The regulation of OPG in liver tissue is largely unknown and we therefore set out to elucidate which growth factors/interleukins associated with fibrosis induce OPG and through which pathways.

**Methods:** Precision-cut liver slices of wild type and STAT6-deficient mice and 3T3 fibroblasts were used to investigate the effects of TGFβ, interleukin (IL) 13 (IL13), IL1β, and platelet-derived growth factor BB (PDGF-BB) on expression of OPG.

**Results:** In addition to TGFβ, only IL13 and not PDGF-BB or IL1β could induce OPG expression in 3T3 fibroblasts and liver slices. This IL13-dependent induction was not shown in liver slices of STAT6-deficient mice and when wild type slices were cotreated with TGFβ receptor 1 kinase inhibitor galunisertib, STAT6 inhibitor AS1517499, or AP1 inhibitor T5224. This suggests that the OPG-inducing effect of IL13 is mediated through IL13 receptor α1-activation and subsequent STAT6-dependent upregulation of IL13 receptor α2, which in turn activates AP1 and induces production of TGFβ and subsequent production of OPG.

**Conclusion:** We have shown that IL13 induces OPG release by liver tissue through a TGFβ-dependent pathway involving both the α1 and the α2 receptor of IL13 and transcription factors STAT6 and AP1. OPG may therefore be a novel target for the treatment liver fibrosis as it is mechanistically linked to two important regulators of fibrosis in liver, namely IL13 and TGFβ1.

## 2. Introduction

Liver fibrosis is a chronic disease induced by long term injury and/or inflammation initiated by virus infections or chemical-induced injury, for example drugs or alcohol [1]. The main pathological characteristic of liver fibrosis is persistent extracellular matrix formation by hepatic stellate cells, which in turn prevents the regrowth of functional hepatocytes [2]. The disease has a high burden as there is no possible therapy to reverse the process when it has fully developed and therefore transplantation is the only option [3].

Transforming growth factor β (TGFβ) has been widely studied for many years as one of the central players in liver fibrosis, but this has not yielded any effective new drugs yet [4, 5]. It is therefore likely that the process of fibrosis development is far more complicated than just the actions of TGFβ alone and that we need to understand the different players and interactions better to develop potential drug candidates.

We recently became interested in the actions of osteoprotegerin (OPG, gene name TNFRSF11B) after finding that OPG is produced in high quantities by (liver) fibroblasts, especially after stimulation with TGFβ and that OPG itself can induce expression of TGFβ, indicating a feed-forward loop [6]. Several clinical studies have shown that higher serum levels of OPG are associated with having liver fibrosis/cirrhosis [7–13]. In addition, OPG serum levels are part of a novel diagnostic score called Coopscore^®^ that has better diagnostic performance than Fibrometer^®^, Fibrotest^®^, Hepascore^®^ and Fibroscan™ in chronic hepatitis C-associated fibrosis [8]. Moreover, in our previous studies, we have demonstrated high hepatic OPG production in liver tissue of patients transplanted for liver cirrhosis and in murine models of liver fibrosis.

Osteoprotegerin is well known for its role in protecting bone matrix degradation [14], but little is known about its function in nonbone tissues. In that respect, its role in vascular calcifications is probably best studied, showing that OPG protects against vascular calcification [15]. This contrasts with its known functional influence in bone metabolism in which it induces calcification of bone [14]. This suggests that OPG has more possible functions unrelated to bone and our previous data show its firm associations with fibrotic processes and TGFβ signaling in (myofibroblasts) [6]. However, little is known about the regulation of OPG production in (liver) fibroblasts by other mediators involved in fibrosis [16]. In this study we therefore aimed to further investigate OPG regulation in the liver by studying the effects of several key fibrosis-related growth factors/interleukins and their downstream signaling pathways. These were interleukin (IL) 1β representing a pro-inflammatory and profibrotic mediator, platelet-derived growth factor BB (PDGF-BB), and IL13, both well-known pro-fibrotic mediators for early and late fibrosis respectively.

## 3. Materials and Methods

### Animals

Male and female wild-type C57BL/6 mice were obtained from Harlan (Horst, The Netherlands) and male STAT6(-/-) C57BL/6 mice were bred in the Institute of Translational Immunology, University Medical Center of the Johannes Gutenberg University Mainz, Germany [17]. Animals were kept in cages with a 12 hour of light/dark cycle and received food and water *ad libitum*.

### Precision-cut liver slices

Murine precision-cut liver slices were prepared as described before by De Graaf et al. (2010) [18]. Slices were treated with 5 ng/mL TGFβ (Peprotech, Rocky Hill, US), 10 ng/mL IL13 (Peprotech), 10 ng/mL IL1β (Peprotech), 10 ng/mL PDGF-BB (Peprotech), 10 mM galunisertib (Selleckchem, Munich, Germany), 21 nM AS1517499 (Axon MedChem, Groningen, The Netherlands), and/or 10 μM T5224 (ApexBio, Houston, US) in triplicate for a total of 48 hours and culture medium was refreshed every 24 hours.

### In vitro cell lines

50,000/well 3T3 murine fibroblasts (The American Type Culture Collection, ATCC^®^ CRL-1658) were cultured in standard medium of Gibco^®^ Dulbecco’s Modified Eagle Medium (Thermo Scientific, Waltham, Massachussets, US) containing 4.5 g/L D-Glucose (Sigma-Aldrich, Missouri, US), 2 mM L-Glutamine (Thermo Scientific, Waltham, Massachussets, US), and 10% of fetal calf serum (Biowest, Nuaillé, France). Cells were starved with medium containing 0.5% serum 24 hours prior to other treatments. Treatments with TGFβ, IL13, IL1β, and PDGF-BB were done at similar concentrations as described for the experiments with slices.

### Generation of tissue lysate

Tissue slices were lysed with extraction buffer containing 25 mM Tris (Sigma-Aldrich, Missouri, US), 10 mM sodium phosphate (Sigma-Aldrich), 150 mM NaCl (Sigma-Aldrich, Missouri, US), 0.1% SDS (Sigma-Aldrich, Missouri, US), 1% Triton-X 100 (Sigma-Aldrich, Missouri, US), and protease inhibitor (Thermo Scientific, Waltham, Massachussets, US) and incubated for 5 minutes at room temperature before snap-freezing and stored at −80°C until analysis.

### Osteoprotegerin analysis

Osteoprotegerin was measured in culture supernatants of cells and slices using a murine OPG DuoSet^®^ ELISA kit (R&D Systems, Minneapolis, US) according to the instructions provided by the manufacturer.

### Messenger RNA analysis

Messenger RNA was isolated from cells or slices (three slices per sample, pooled, homogenized prior to extraction) using Maxwell^®^ LEV Simply RNA Cells/Tissue kit (Promega, Madison, Wisconsin, US). A NanoDrop^®^ ND-1000 Spectrophotometer (Thermo Scientific) was used to measure total mRNA concentration in samples. cDNA synthesis from the mRNA was performed using a Moloney Murine Leukemia Virus Reverse Transcriptase (M-MLV RT) kit (Promega, Madison, Wisconsin, USA) in a Mastercycler^®^ Gradient (Eppendorf, Hamburg, Germany) programmed for 10 minutes at 20°C, 30 minutes at 42°C, 12 minutes at 20°C, 5 minutes at 990C, and 5 minutes at 20°C. Transforming growth factor beta 1 (TGFβ1), IL13 receptor α2 (IL13Rα2), pro-collagen 1 subunit α1 (Col1α1), α-smooth muscle actin (αSMA), heat shock protein 47 (HSP47), plasminogen activator inhibitor 1 (PAI1), and fibronectin 1 (Fn1) genes were quantified using quantitative real time PCR (RT qPCR) from the synthesized cDNA, using SensiMixTM SYBR^®^ Green (Bioline, London, UK) in a 7900HT Real-Time PCR sequence detection system (Applied Biosystems, Waltham, Massachussets, US) with primer sequences as presented in Table 1. PCR analysis consisted of 45 cycles of 10 min at 95°C, 15 seconds at 95°C, and 25 seconds at 60°C (repeated for 40 times) followed by a dissociation stage of 95°C for 15 seconds, 60°C for 15 seconds, and 95°C for 15 seconds. Output data were analyzed using SDS 2.4 software (Applied Biosystems) and ΔCt values were calculated after β-actin normalization. Two to the power of −ΔCt (2^-ΔCt^) was used as a final value to be statistically analyzed.

**Table 1.**
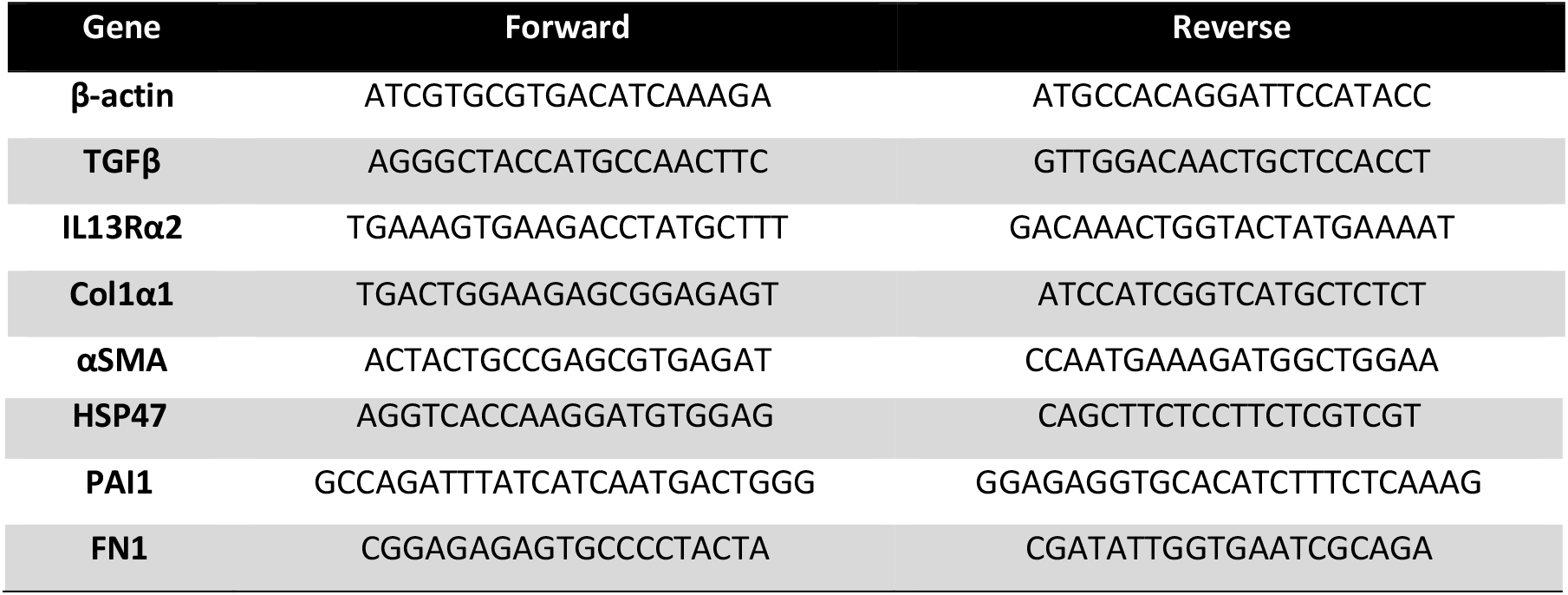
Primers sequences I would expand this legend a bit. At least mention human/mouse

### Viability assay

Viability of the slices was assessed by measuring the ATP content per milligram tissue using a bioluminescence assay kit (Sigma-Aldrich) as previously reported by Hadi et al. [19]. For each sample, three slices were collected separately in 1 mL sonification optimization (SONOP) solution pH 10.9 containing 70% ethanol and 2 nM EDTA.

### Statistics

All statistics were performed using GraphPad Prism 8. As datasets were all n<8, nonparametric tests were used. When comparing 2 groups a Mann Whitney U or Wilcoxon test was used depending on the data being paired or not. When comparing multiple groups, a Friedman or Kruskall-Wallis with Dunn’s correction was used. Data are presented as min-to-max box-and-whisker plots with individual data points. For the time course experiment using 3T3 fibroblasts, the areas under the curve from 0.5-12 hours and 12-36 hours were calculated and these were compared between groups. Data in this experiment are presented a median + the interquartile range. For all experiments, p<0.05 was considered significant.

## 4. Results

### IL13 induces fibroblast and hepatic OPG production

To study possible factors that can induce OPG production by fibroblasts, we treated 3T3 fibroblasts with several cytokines associated with fibrosis. In this study we used a major pro-inflammatory and profibrotic cytokine IL1β, and pro-fibrotic cytokines IL13 and PDGF-BB with TGFβ as our positive control as we have shown higher OPG expression with TGFβ in our previous study [6]. In addition to TGFβ, only IL13 resulted in higher OPG production as compared to control (figure 1A). To confirm that IL13 can have a similar effect in liver tissue, we treated murine precision-cut liver slices with IL13 using TGFβ again as a positive control and found that IL-13 also resulted in significantly higher OPG release from liver tissue as compared to control (figure 1B). This higher OPG release in slices was accompanied by near-significant higher OPG mRNA expression and significant higher expression of fibrosis-associated genes col1α1, HSP47, and FN1, but not αSMA and PAI1 (figure 1C). None of the treatments affected the viability of the slices (supplemental figure S1).

**FIGURE 1.**
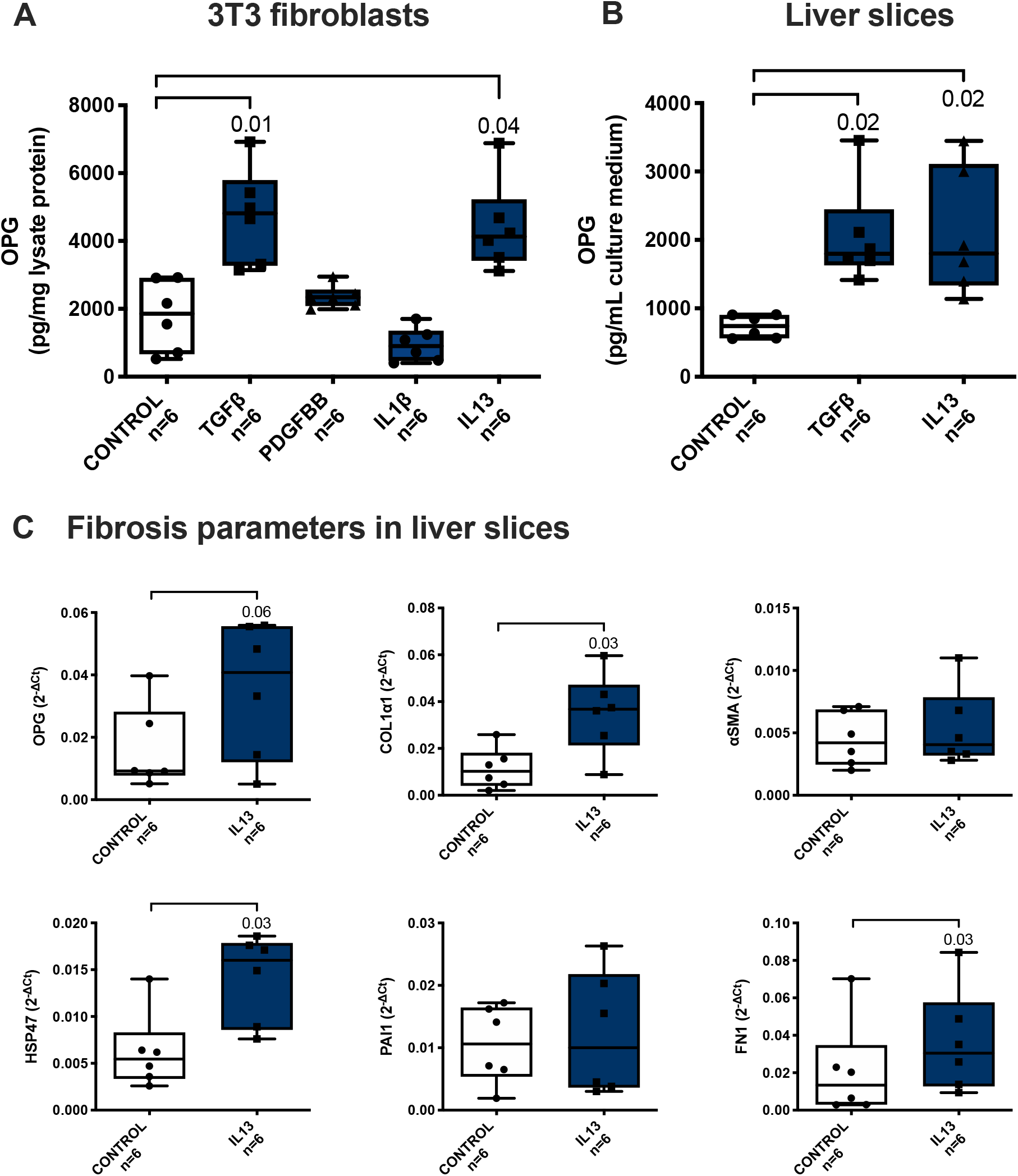
IL13 induces OPG release. When 3T3 fibroblasts were treated with IL1β, IL13, PDGFBB, or TGFβ1 (positive control) for 24 hours of incubation, only IL13 and TGFβ1 treatment resulted in significantly higher OPG release as compared to control (**A**). Murine precision-cut liver slices also released significantly more OPG in culture medium after 48 hours of incubation with IL13 as compared to control, just like positive control TGFβ1 (**B**). IL13 incubation also resulted in (near)significant higher expression of the fibrosis-associated genes OPG, Col1α1, HSP47, and FN1, though not of αSMA and PAI1 (**C**). Groups were compared using a Friedman test with Dunn’s correction or a Wilcoxon test and p<0.05 was considered significant.

### IL13 induces OPG production at a slower rate than TGFβ

To check whether induction of OPG production followed similar kinetics between TGFβ and IL13, we followed OPG release in time in culture medium of 3T3 fibroblasts after stimulation with TGFβ and IL13. We found that after 36 hours of incubation IL13 and TGFβ both induced a similar release in OPG although the induction by TGFβ occurred somewhat faster. When comparing the area under the curve between stimulated cells and untreated control cells in the first 12 hours, we found a significant increase in OPG release by TGFβ, while IL13 was not significantly different from control. In the time interval from 12 to 36 hours both TGFβ and IL13 significantly induced OPG release as compared to control (figure 2).

**FIGURE 2.**
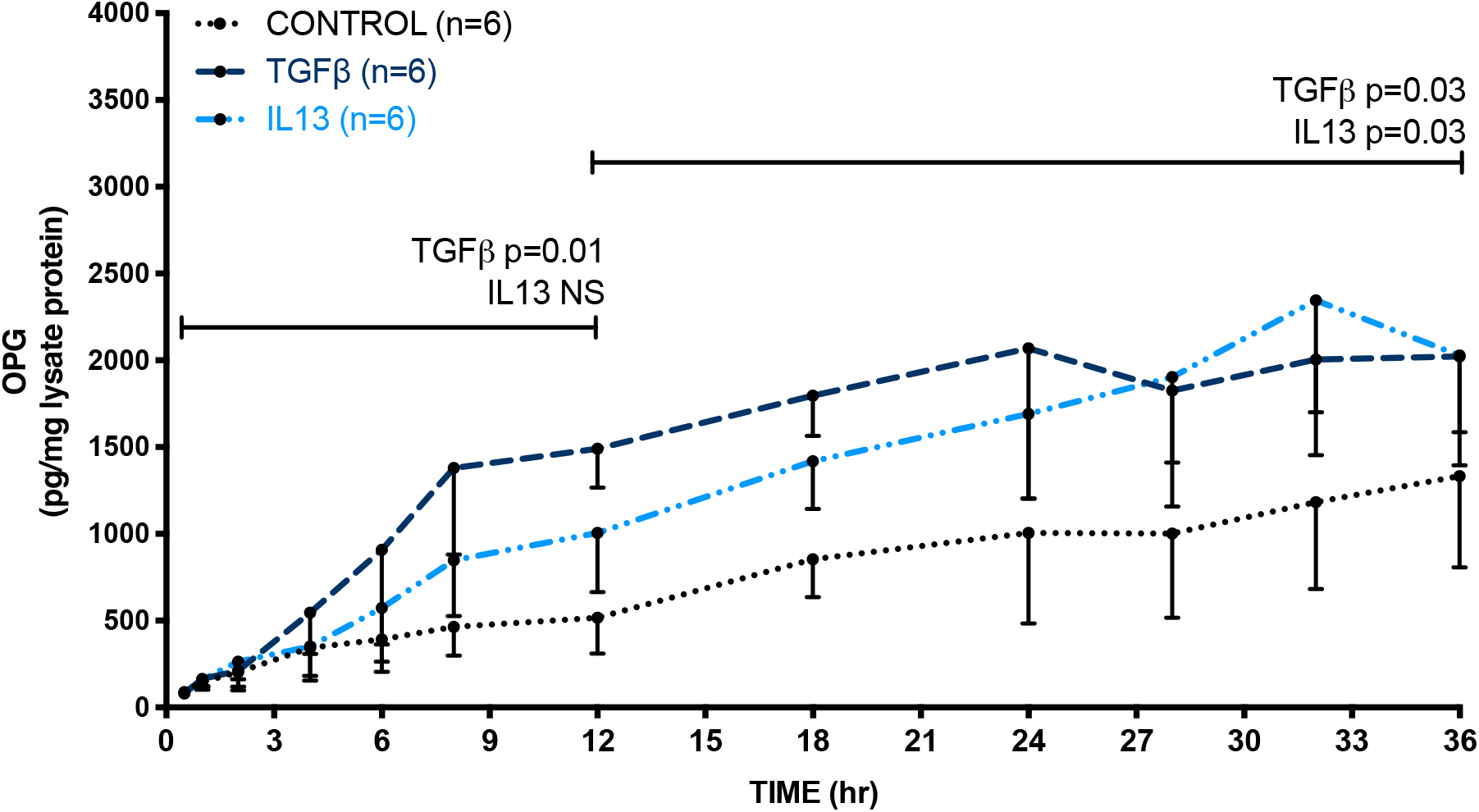
IL13 induces OPG at a slower rate than TGFβ. 3T3 cells were incubated with TGFβ or IL13 and OPG release in medium was measured at several time points up to 36 hours. TGFβ was shown to significantly upregulate OPG release already in the first 12 hours as compared to control, whereas IL13 needed more time for a similar effect. Groups were compared using a Friedman test with Dunn’s correction and p<0.05 was considered significant.

### IL13 induces hepatic OPG induction through TGFβ

We hypothesized that TGFβ may be involved in the higher hepatic OPG production by mouse liver tissue after IL13 treatment as IL13 has been shown to induce TGFβ1 expression [20]. We therefore assessed TGFβ1 mRNA expression in liver slices after incubation with IL13 and we found a trend towards higher TGFβ1 mRNA expression after IL13 treatment compared to untreated control slices (figure 3a). To confirm that TGFβ is indeed involved in the IL13 effect on OPG induction, we also incubated liver slices with galunisertib, a TGFβ1 receptor inhibitor, together with IL13. We found that with galunisertib cotreatment, IL13 treatment did not result in higher OPG release from liver tissue anymore (figure 3b). None of the treatments affected the viability of the slices (supplementary figure S1).

**FIGURE 3.**
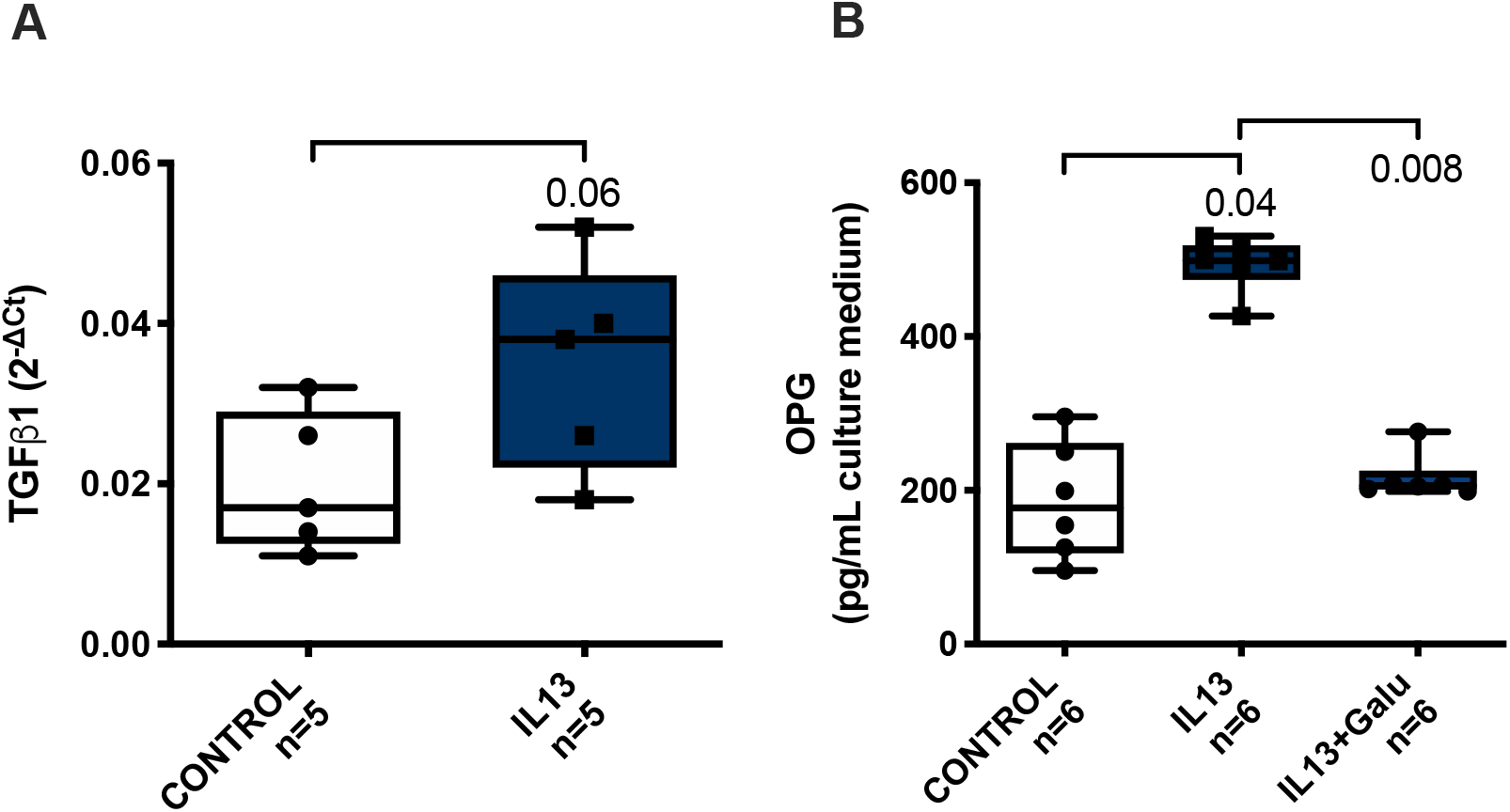
IL13 induces OPG through TGFβ1. When mouse liver slices were treated with IL13, we found a trend towards more TGFβ1 mRNA expression. Groups were compared with a Wilcoxon test, p<0.05 was considered significant (**A**). Moreover, when IL13 was given together with galunisertib, a TGFβ1-receptor inhibitor, the IL13-induced higher OPG release was not found anymore. Groups were compared with a Friedman test with Dunn’s correction, p<0.05 was considered significant (**B**).

### STAT6 is involved in ILl3-induced release of OPG

IL13 has been reported to signal through 2 receptors: receptor IL13Rα1 and IL13Rα2 [21]. The downstream activation pathway of IL13Rα1 is via transcription factor STAT6 [22]. To study whether the activation of IL13Rα1 and subsequently STAT6 is involved in the IL13-induced release of OPG, we treated liver slices of STAT6-deficient mouse with IL13 or TGFβ and measured OPG released in medium. We found that IL13 failed to induce OPG release by liver slices of STAT6-deficient mice as compared to untreated controls, whereas TGFβ could still induce OPG release as we found before in wildtype mice (figure 4a). To confirm our finding, we used AS1517499, a chemical compound blocking STAT6 activity, in our wild-type mouse liver slices [23] and similarly found that IL13 did not induce OPG release anymore when slices were co-incubated with this inhibitor as compared to slices only treated with IL13 (figure 4b). None of the treatments affected the viability of the slices (supplementary figure S1).

**FIGURE 4.**
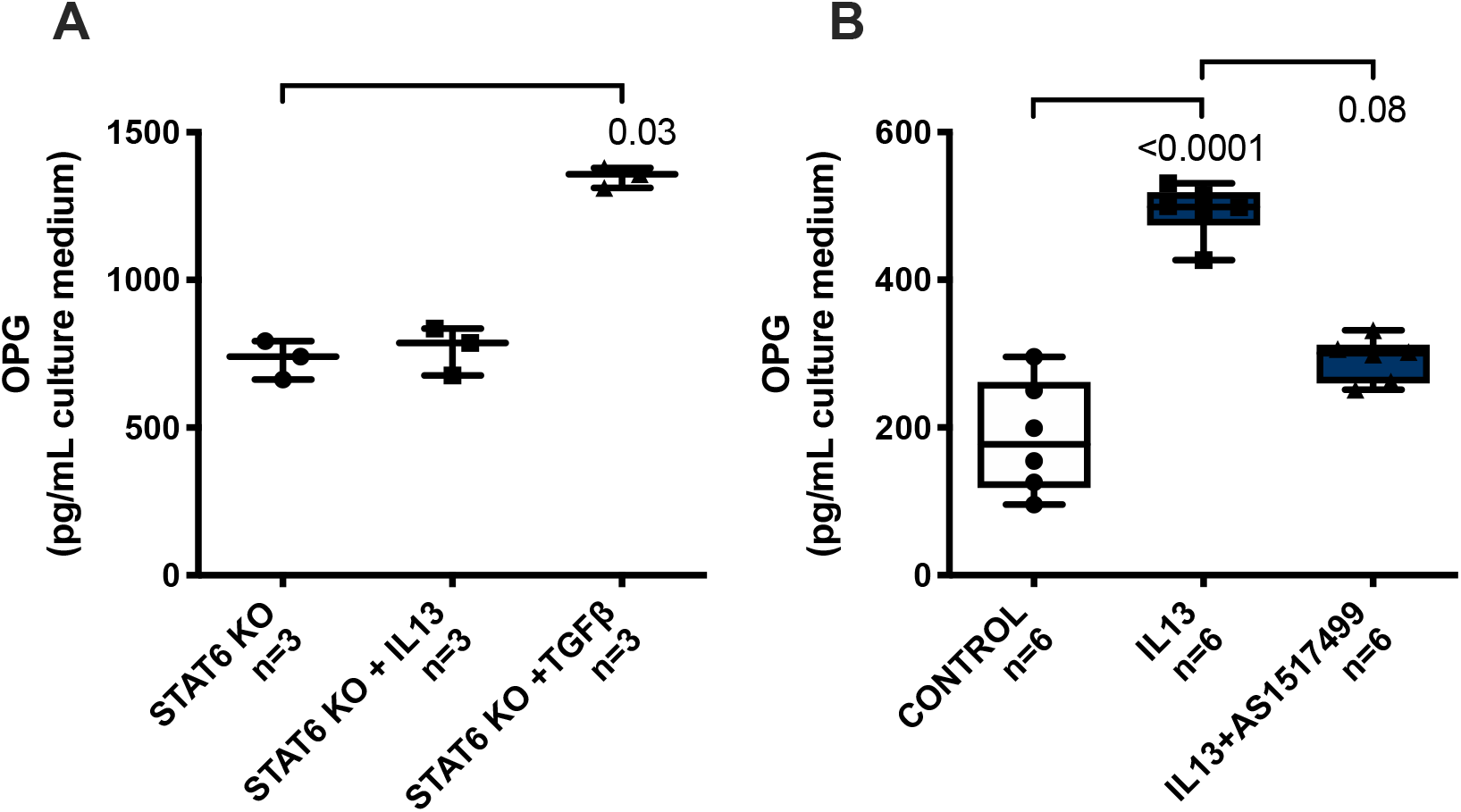
STAT6 is involved in IL13-induced release of OPG. When liver slices of the STAT6 knock out (KO) mice were incubated for 24 hours with IL13 or TGFβ1, IL13 failed to induce OPG release, while the TGFβ1-induced release was not affected by the deficiency in STAT6. Groups were compared using a Friedman test with Dunn’s correction, p<0.05 was considered significant (**A**). IL13-induced OPG release in wild type liver slices could also be blocked with AS1517499, an inhibitor of STAT6 activity. Groups were compared using a Friedman test with Dunn’s correction, p<0.05 was considered significant (**B**).

### IL13 receptor α2 is also involved in IL13-induced OPG release

Fichtner-Feigl et al. (2005) reported that IL13Rα2 is involved in induction of TGFβ expression and fibrosis through transcription factor AP1 [24]. However, in homeostatic conditions, the expression of this receptor is low [25], while activation of IL13Rα1 and subsequently STAT6 can induce IL13Rα2 expression [26]. In order to check whether these findings are also relevant in our system, we assessed IL13Rα2 mRNA expression in liver slices upon IL13 treatment. We found that IL13Rα2 mRNA expression level was significantly higher upon IL13 treatment as compared to untreated controls (figure 5a). We then used T5224, a chemical inhibitor of AP1 [27] to study whether AP1 in involved in IL13-induced OPG release and we found that indeed chemical inhibition of AP1 completely abolished the IL13-induced release of OPG (figure 5b). None of the treatments affected the viability of the slices (supplementary figure S1).

**FIGURE 5.**
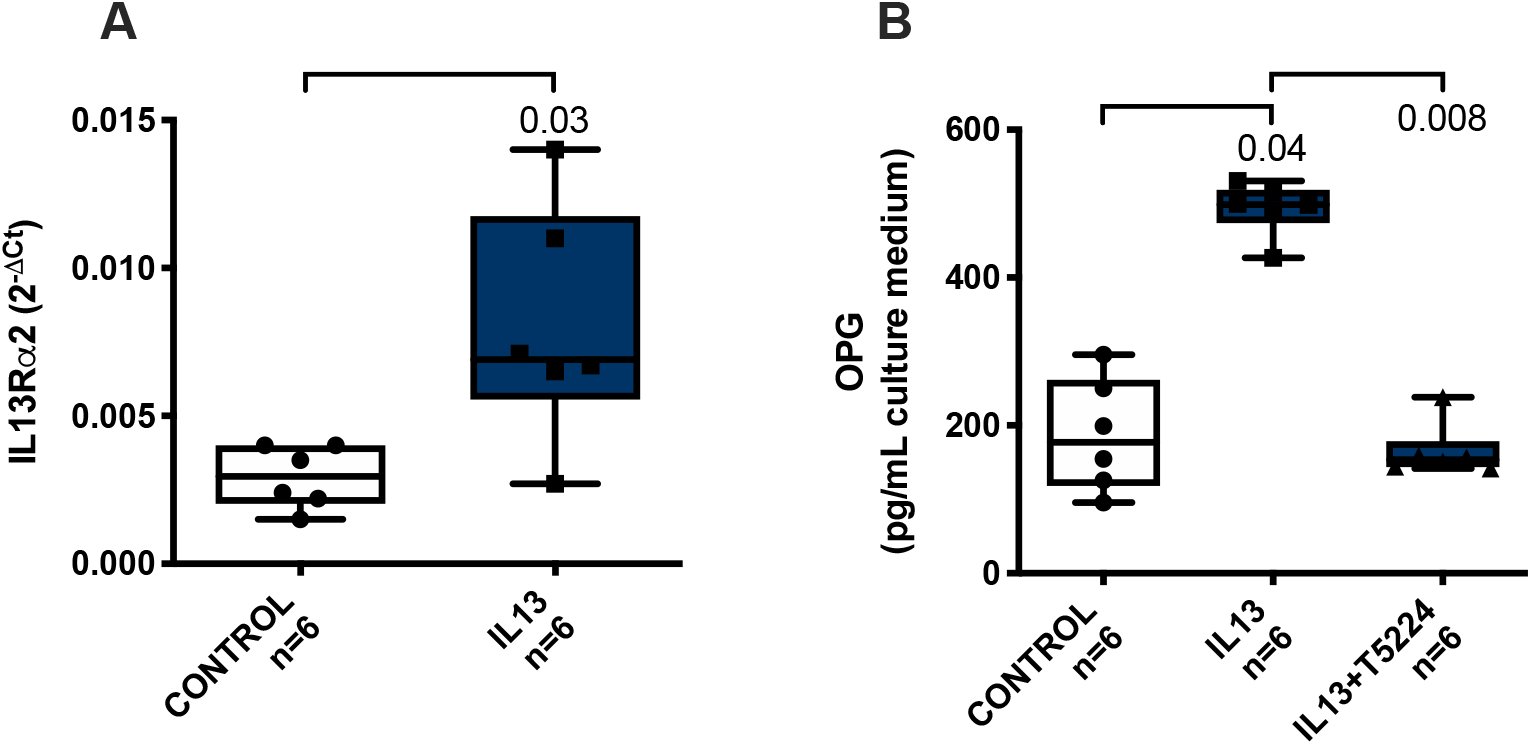
IL13 receptor α2 is also involved in IL13-induced OPG release. Mouse liver slices treated with IL13 had higher IL13Rα2 mRNA expression as compared to untreated controls. Groups were compared using a Wilcoxon test, p<0.05 was considered significant (**A**). Mouse liver slices cotreated with IL13 and T5224, an AP1 inhibitor, did not show higher OPG release as compared to IL13 treatment alone. Groups were compared using a Friedman test with Dunn’s correction, p<0.05 was considered significant (**B**).

## 5. Discussion

We have previously shown higher OPG expression in fibrotic/cirrhotic conditions and that TGFβ can induce this hepatic OPG production [6]. Moreover, OPG in its turn was shown to induce TGFβ expression, contributing to a feed-forward loop. This study now shows that in addition to TGFβ, also the fibrosis-associated cytokine IL13, known to induce collagen expression and promote liver fibrosis via stat-6 signaling [28, 29], can induce OPG expression and release by murine liver tissue. Interestingly, this IL13-induced OPG production is completely dependent on TGFβ through a pathway involving IL13Rα1/STAT6 and IL13Rα2/AP1. The strength of this study is that we used precision-cut liver tissue slices, instead of cultures of individual cells or cell lines, making our results more relevant for in vivo situations.

In our previous studies we identified fibroblasts as the main source of OPG production in liver during fibrogenesis and after TGFβ exposure. Little is known about the regulation of OPG production in these cells and in liver tissue itself. We therefore investigated the effect on OPG production of other cytokines/growth factors involved in fibrogenesis. For this we chose IL13 and PDGF-BB as these were identified as important fibrogenic regulators of fibroblasts and therefore fibrosis, and are being investigated as potential targets of antifibrotic drugs [30–32]. We also chose IL1β because this key cytokine is involved in chronic liver inflammation and subsequent development of fibrosis [33]. Our results showed no influence of IL1β on the production of OPG by fibroblasts. This finding suggests that OPG is mostly produced in connection to fibrogenesis and not during inflammation that may precede the development of fibrosis. Furthermore, PDGF-BB did not induce OPG production in fibroblasts either. PDGF-BB is the mitogenic agent for fibroblasts and triggers proliferation and migration of these cells. The pathways leading to stimulation of proliferation by PDGF-BB are apparently not linked to stimulation of OPG production.

IL13 stimulation on the other hand, did lead to higher production of OPG in fibroblasts. This effect was confirmed using precision-cut liver slices of murine livers and this experiment also showed us that the higher OPG expression and production after IL13 stimulation was accompanied by higher expression of fibrosis-associated markers Col1α1, HSP47, and Fn2. These profibrotic results of IL13 are in line with previously published results by Sugimoto et al. (2005) [34] and Gieseck et al. (2016) [30], who showed that IL13 can induce collagen production and fibrogenesis in hepatic stellate cells and liver tissue respectively. Induction of fibrogenesis by IL13 has been suggested to occur via upregulation of TGFβ1 via IL13Rα1 and IL13Rα2 signaling [24, 34–36], although other studies have suggested that IL13 can also induce fibrosis independently from TGFβ [37]. Our data show that the IL13-induced production of OPG is completely dependent on TGFβ1, as inhibition of TGFβ1-signalling with galunisertib completely abrogated the effect of IL13 on OPG production. We also confirmed that both IL13 receptors are involved in this TGFβ1-mediated OPG production in liver tissue [24, 34–36]. Blocking or the absence of STAT6 fully blocked IL13-induced OPG production by liver tissue, showing that IL13Rα1-signalling through STAT6 is necessary to upregulate TGFβ1 and subsequently OPG. Furthermore, we showed that IL13 stimulation leads to higher expression of IL13Rα2 in liver tissue and that blocking signalling of this receptor with an AP1 inhibitor also completely abrogated IL13-induced OPG production in liver tissue. A scheme explaining these main findings is depicted in figure 6.

**FIGURE 6.**
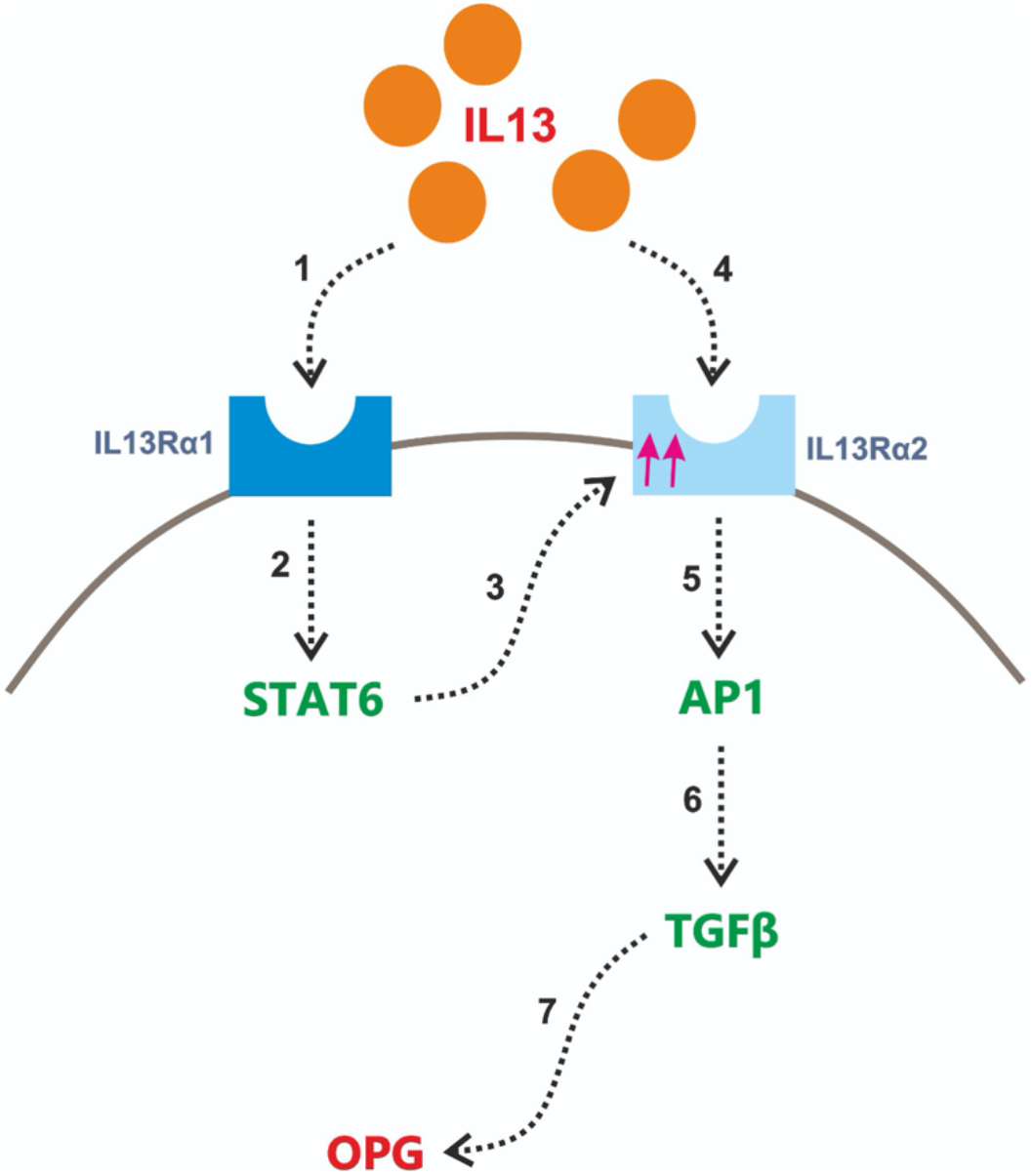
IL13 induces liver OPG production via activation of both IL13 receptors and subsequent induction of TGFβ1. A schematic overview of IL13-induced OPG production based on the results presented in this study. (1) IL13 binds to receptor α1 followed by activation of this receptor triggering (2) STAT6 activation resulting in (3) the increased expression of IL13 receptor α2, which is initially expressed at low levels. (4) IL13 then binds to receptor α2, triggering (5) activation of transcription factor AP1, which induces (6) expression of TGFβ1. Finally, as we have reported in our previous study [6], TGFβ1 can induce OPG protein production by the liver (7).

The signaling through both IL13 receptors and the need for TGFβ1 upregulation before OPG can be produced, probably also explains why IL13-induced OPG production was lagging in comparison to TGFβ1-induced OPG production. In 36 hours, both cytokines can induce production of similar levels of OPG, but we found that TGFβ1 can achieve these levels faster. IL13Rα2 is expressed at low levels in normal liver tissue and can therefore not induce TGFβ1 expression immediately when liver tissue is exposed to IL13. The fact that IL13 first needs to upregulate IL13Rα2 through IL13Rα1 and STAT6 to be able to stimulate TGFβ1 expression probably accounts for the delayed production of OPG after IL13 stimulation.

Our results point at an interesting feed-forward mechanism in liver tissue involving TGFβ1 as a central player, with IL13 prolonging profibrogenic TGFβ signaling via induction of the TGFβ inducer OPG. Both TGFβ1 and IL13 can induce OPG and OPG can induce expression of TGFβ1 again [6]. In most cases, liver tissue damage will result in normal repair and restoration of fully functional liver tissue. Therefore, there must also be brakes in this process to prevent that any type of damage will always end in fibrosis. Our current studies focus on several microRNAs that are induced by TGFβ1 and that may serve as the brakes in this feed-forward loop. Another interesting aspect of our findings is the possibility of using OPG as a target for therapy. Both TGFβ1 and IL13 are targets for inhibition of fibrogenesis that are currently being explored in clinical trials [37–41]. As OPG is linking both pathways it seems to be another promising target for therapy.

## 6. Conclusion

We have shown that IL13 induces OPG release by liver tissue through a TGFβ-dependent pathway involving both the α1 and the α2 receptor of IL13 and transcription factors STAT6 and AP1. OPG may therefore be a novel target for the treatment liver fibrosis as it is mechanistically linked to two important regulators of fibrosis in liver, namely IL13 and TGFβ1.

## 7. Supplementary Material

Treatments for 48 hours of compounds in our experiments did not significantly compromise the viability of the mouse liver slices used in our study (n=6, Kruskal-Wallis test corrected for multiple testing).

## 8.1. Acknowledgement

DS received project-related support from EU Horizon 2020 projects under grant agreements nr. 634413 (EPoS, European Project on Steatohepatitis).

## 8.2. Statement of Ethics

The use of C57BL/6 mice in this study was approved by the Institutional Animal Care and Use Committee of the University of Groningen (DEC 6416 AA) and the use of STAT6(-/-) mice by the Institutional Animal Care and Use Committee of the Government of Rhineland Palatinate under the reference number 2317707/G12-1-007.

## 8.5. Author Contributions

Conceptualization, A.A., L.B., and B.N.M., data curation, A.A., K.S.S.P., E.G., K.A.M., C.R.-S., and B.N.M., formal analysis: A.A., L.B., and B.N.M., funding acquisition, P.O., and B.N.M., investigation, A.A., L.B., and B.N.M., methodology, A.A., K.S.S.P., L.B., C.R.-S., D.S., P.O., and B.N.M., project administration, B.N.M., resources, B.N.M., software, B.N.M., supervision, L.B., P.O., and B.N.M., validation, L.B. and B.N.M., visualization, A.A., L.B., and B.N.M., writing–original draft, A.A., K.S.S.P. and B.N.M., writing–review and editing, K.A.M., C.R.-S., D.S., L.B., P.O., and B.N.M. All authors have read and agreed to the published version of the manuscript.

## 11. Supplementary Figure and Legend

**Supplementary Figure S1.**
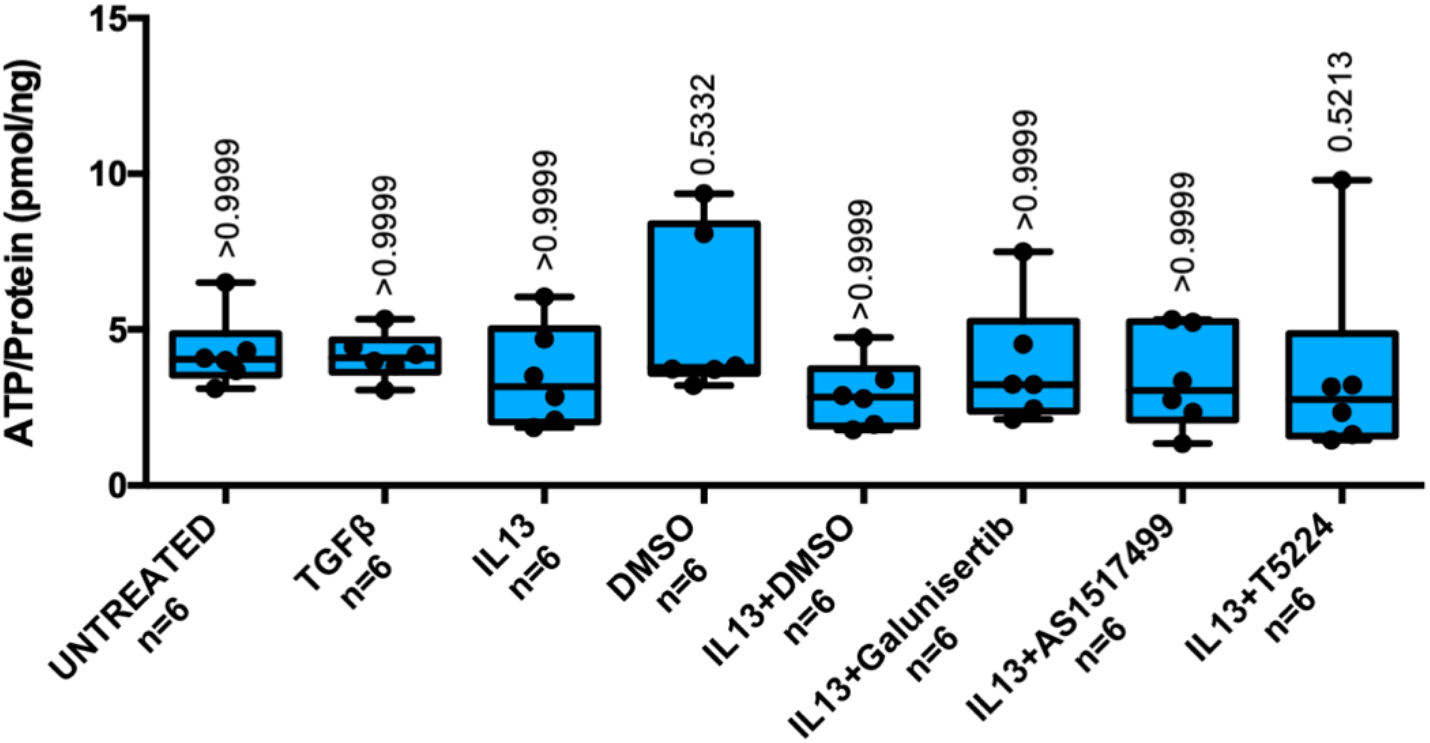
Treatments did not compromise viability mouse liver slices. Treatments for 48 hours of compounds in our experiments did not significantly compromise the viability of the mouse liver slices used in our study (n=6). Groups were compared using a Kruskal-Wallis test corrected for multiple testing, p<0.05 was considered significant.

